# Pathogenic *Leptospira* species identified in dogs and cats during neutering in Thailand

**DOI:** 10.1101/2025.08.01.668072

**Authors:** Metawee Thongdee, Somjit Chaiwattanarungruengpaisan, Weena Paungpin, Sivapong Sungpradit, Sineenard Jiemtaweeboon, Ekasit Tiyanun, Kanin Ruchisereekul, Sarin Suwanpakdee, Janjira Thaipadungpanit

## Abstract

Pathogenic species of Genus *Leptospira* cause an underdiagnosed zoonosis in humans and animals called leptospirosis. Animal reservoirs getting infections without signs can carry and shed the active spirochete into the environment through their urine, making the control of leptospirosis transmission to humans more challenging. Asymptomatic leptospirosis in human companions, such as dogs and cats, leading to unawareness of infections, has been demonstrated in a few countries. Monitoring disease prevalence and investigating molecular epidemiology is essential for controlling and preventing human and animal leptospirosis transmission.

To investigate the prevalence of *Leptospira* infection among cats and dogs in Thailand, we collected 567 urine samples through a neutering program in seven provinces across the central, western and southern regions. The urine samples were screened for *Leptospira*’s DNA and further speciation using the Sanger Sequencing Analysis. 34/303 (11.2%) dogs and 22/264 (8.3%) cats tested positive for *Leptospira* in the Pathogen clade. The partial *rrs* gene analysis identified *L. interrogans*, *L. weilii*, and *L. borgpetersenii* (4.4%). We first reported *L. yasudae* (1.0%) and other new species of the Pathogen subclade 2 (1.4%) infections among these dogs and cats.

Three key pathogenic *Leptospira* species commonly causing human leptospirosis in Southeast Asia were identified in healthy owned and free-roaming dogs and cats, suggesting the risk of human leptospirosis in the areas. These animal reservoirs (probably living under the same roof as us) can transmit diseases daily to owners (caretakers), other animals and the environment. Effective public awareness campaigns on the transmission might encourage dog and cat health surveillance and vaccinations, reducing human infection risks.

**Author Summary:** Leptospirosis, a neglected zoonotic disease caused by pathogenic species of genus *Leptospira*, can be transmitted to humans through contact with infected animal body fluids or contaminated environments. Free-ranging dogs and cats can shed infectious *Leptospira* in their urine, posing a significant risk. In Thailand, animal vaccine coverage for diseases like leptospirosis in these animals is unknown and unregulated, raising concerns about zoonotic transmission and reflecting a lack of routine veterinary support.

Our study joined non-profit, volunteer-led neutering programs that travel to various provinces, offering free services for owned and unowned pets. This allowed our team to collect urine specimens directly from the urinary bladder for the detection of *Leptospira*. We found nearly 10% of the recruited asymptomatic dogs and cats were infected by the molecular screening assay. Three pathogenic *Leptospira* species, commonly infecting humans globally, were identified. Notably, we report the first detection of *L. yasudae* in animal urine samples, providing evidence of infectivity for a species not previously recognised for its pathogenicity and typically reported from the environment. This highlights the risk of human leptospirosis from contact with cats and dogs, emphasising the need for public awareness and annual vaccination programs for all pets.

## Introduction

The genus *Leptospira* is a gram-negative spirochete bacterium. The primary pathogenic species that cause human leptospirosis worldwide are *L. interrogans* and *L. borgpetersenii* [1, 2]. Many mammals, including rodents, dogs, cats, pigs, bats, and cattle, can carry these spirochetes in their proximal renal tubules without clinical signs. They excrete living bacteria in their urine as reservoir hosts [3, 4]. Direct exposure to infected animal urine or indirect exposure to a contaminated environment contributes to disease transmission [3–5]. Disease severity depends on bacterial virulence, inoculum size and host immunity [5].

Of 68 species in the genus *Leptospira*, 40 are categorised into the Pathogen (P) clade, and 28 are in the Saprophyte (S) clade based on whole genome sequence analysis [6–12]. The P clade comprises two subclades (P1 and P2). Of 19 species in subclade P1, eight (*L. interrogans*, *L. kirschneri*, *L. noguchii*, *L. alexanderi*, *L. weilii*, *L. borgpetersenii*, *L. santarosai*, and *L. mayottensis*) are key species that cause leptospirosis worldwide. The other eleven species (*L. adleri, L. ainazelensis, L. ainlahdjerensis, L. alstonii, L. barantonii, L. ellisii, L. gomenesis, L. kmetyi, L. stimsonii, L. tipperaryensis,* and *L. yasudae*) are recently discovered species isolated from environmental samples, with no evidence of human or mammalian infections. Of 21 species of the P2, five (*L. inadai*, *L. fainei*, *L. broomii*, *L. wolffii*, and *L. licerasiae*) have been known as the intermediate group, causing mild leptospirosis in mammals and humans [11, 13–17].

Human leptospirosis was commonly reported in tropical and subtropical regions [18, 19]. Over one million global human infections and more than 58,000 fetal cases were reported between 1970 and 2008 [18]. In Thailand, 24,378 human patients were reported to the Bureau of Epidemiology, Ministry of Public Health between 2017 and 2021 (average incidence rate of 3.6 per 100,000 population per year [20]. The incidence rates in the lower northeastern (11.5 per 100,000 population per year) and upper southern (9.7 per 100,000 population per year) regions were higher than those in other regions of Thailand. Based on the computational models, increasing rainfall, flood exposure, using multiple household water sources, and occupational exposure to soil with high moisture, environmental water, and animals affected leptospirosis transmission among animals, environments, and humans [21–23].

Leptospirosis in animals can pose a public health threat, as asymptomatic infected animals can spread living bacteria through urination, contaminating habitats and foraging areas where humans live, travel, or work. The more exposure to a variety of probable infection sources, the higher the risk of infections, which shows the complexity of disease control [24, 25]. Pets can transmit disease to their owners by bringing pathogens from outside into their houses. Dogs and cats are common pets globally, including in Thailand, as over one-third of households own at least one cat or dog [26]. Of more than 12 million cats and dogs in Thailand, estimated between 2019 and 2020, one million were free-roaming [27–29]. *Leptospira* carriage was identified in free-roaming or client-owned dogs and cats, and its frequency varied among geographical locations [30–32].

Symptomatic and asymptomatic leptospirosis were reported worldwide in up to 97% of dogs and 68% of cats since 2003 [30–70]. The infections caused by the following *Leptospira* species in the P1 were commonly reported. *L. interrogans* and *L. borgpetersenii* were found in dogs (0.4-16% vs 0.5-1%) [25, 31, 33, 34, 36, 37, 41, 42, 47–51, 55, 57, 69, 71] and cats (2-5% vs 0.7-7%) [58, 61–64, 70] while *L. weilii* (1-3%) [31, 33], *L. santarosai* (0.5-37%) [42, 48, 54, 57], *L. noguchii* (2-37%) [25, 50], *L. kirschneri* (0.4%) [69] and *L. kmetyi* (3-10%) [33, 48] were found in dogs only. In comparison, the P2 (*L. wolffii* and *L. licerasiae*) infections were reported in 18% and 5% of dogs originating from Iran and Sri Lanka, respectively [48, 71]. Cross-sectional investigations of asymptomatic *Leptospira* infections in dogs and cats in Thailand are scarce [31, 32]. In this study, we aimed to investigate *Leptospira* infections in dogs and cats residing in three regions of Thailand, providing epidemiological data to inform the development of leptospirosis transmission prevention and control policies for dogs and cats.

## Methods

### Ethics statement

The collection and analysis of animal samples were conducted between 2020 and 2023 under the approval of the Institute for Animal Care and Use Committee, Faculty of Veterinary Science, Mahidol University (MUVS-2020-01-03). Written informed consent was obtained from animal owners for their animals’ participation. The unowned or free-roaming dogs and cats primarily resided in public areas frequented by local communities, such as temples and schools. Informed written consent for a sample collection was obtained from the heads of each community. The laboratory protocols were approved by the Institutional Biosafety Committee of the Faculty of Veterinary Science, Mahidol University (IBC/MUVS-B-001/2564).

### Sample size calculation

Sample size estimation was performed using ProMESA software version 2.3 (EpiCentre, Massey University, New Zealand) based on the previously reported leptospirosis prevalence at 10.3% [31] of dogs and 0.8% of cats [32] at a 95% confidence interval and a 2% relative error. This resulted in a minimum sample size of 223 dogs and 236 cats, which was required for the study.

### Study population

Dogs and cats that look healthy underwent sterilisation through the neutering program of the Faculty of Veterinary Science, Mahidol University and nonprofit organisations. All owned and not-owned (free-roaming) dogs and cats were included in this study. The neutering program was carried out in seven provinces of three regions in Thailand: central (Nakhon Sawan, Nakhon Pathom and Samut Sakhon provinces), western (Tak, Kanchanaburi and Prachuap Khiri Khan provinces) and southern (Ranong province) (Fig 1). The sex and estimated age of each animal were recorded.

**Fig 1.**
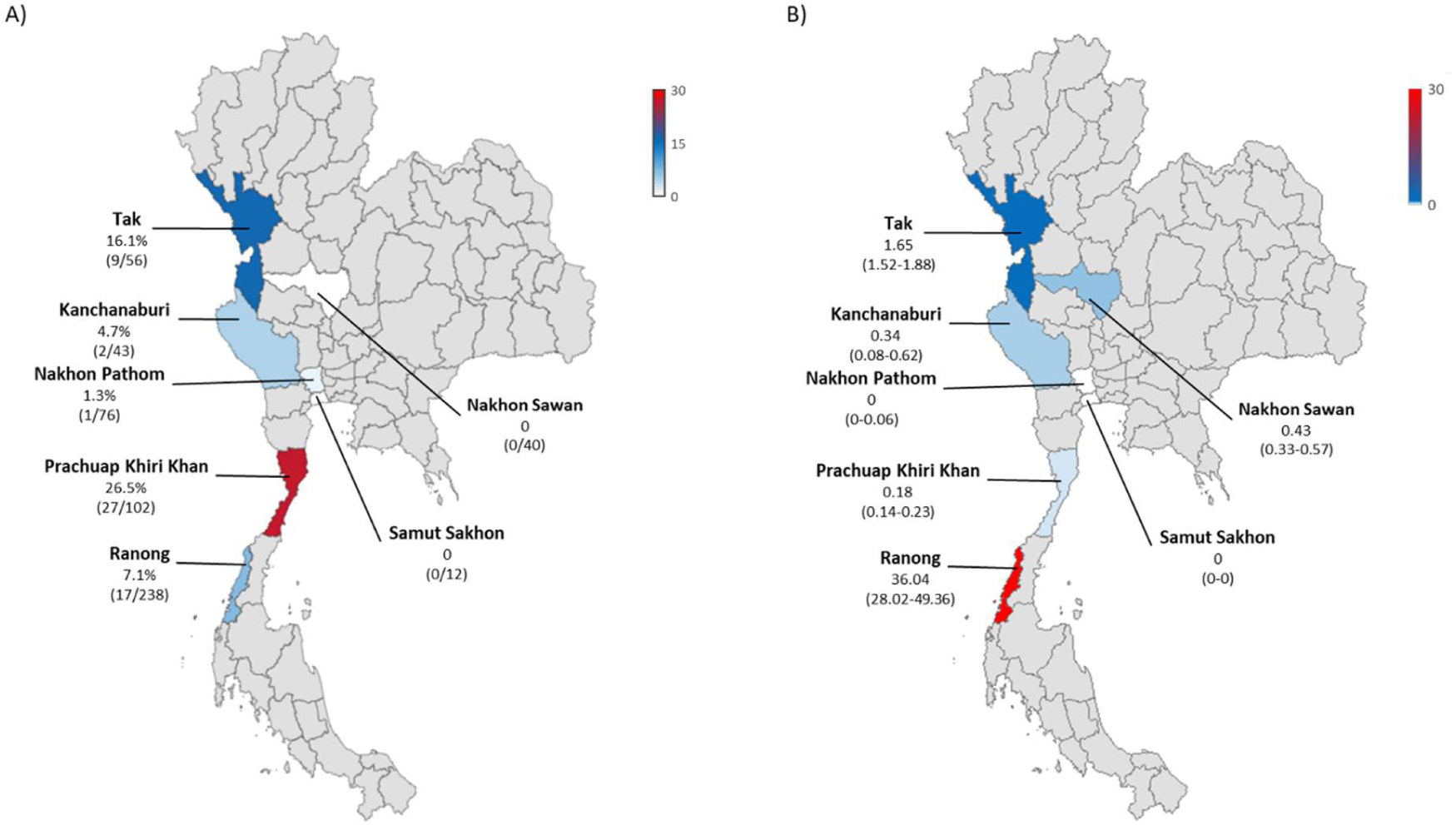
The Thailand maps show the geographical distributions of A) dog and cat *Leptospira* infections identified using PCR assays and partial 16S rRNA gene sequencing analysis, compared to B) those of the median annual incidence of human leptospirosis reported to the Department of Disease Control, Ministry of Public Health, Thailand. The provinces of the seven study sites, located in three regions (central, western, and southern), are highlighted. The darker the colour, the higher the infection frequency or incidence. While the hotter the colour, the higher the infection frequency or incidence. In panel A, the infection frequencies are shown, along with the proportions of infected animals among the total number of dogs and cats investigated in the study. Panel B displays the median annual incidence of human leptospirosis per 100,000 population from 2019 to 2022, along with the interquartile range in parentheses. These reported cases were diagnosed based on clinical symptoms alone or with serology-based or PCR-based assays.

### Dog and cat urine collections and DNA preparations

Urine samples were collected during trips with the non-profit, volunteer-led neutering programs. These programs provided access to anesthetised animals from which urine samples were collected using sterilised urinary catheters before sterilisation surgery. Urine samples were kept on ice upon collection until DNA preparation was completed in laboratories near the study sites. Collected urine specimens’ volume ranged from 1-370 ml (median = 42 ml) in dogs and 0.8–108 ml (median = 9.5 ml) in cats. The total volume of each collected sample contained in a sterile 50 ml tube was centrifuged at 2,000 relative centrifugal force (RCF) for 10 minutes at 4°C to remove cells and debris. Then, the supernatant was transferred to a new sterile 50 ml tube and centrifuged again at 20,000 RCF for 30 minutes at 4 °C to obtain a microorganism cell pellet for the DNA preparation using the Genomic DNA Mini Kit (blood and cultured cells) (Geneaid, New Taipei City, Taiwan) according to the manufacturer’s protocol. The extracted DNA samples were eluted using 30-40 μl of the elution buffer.

### Identification of *Leptospira* species based on partial 16S rRNA sequences amplification and analysis

The nested PCR assay targeting the partial 16S rRNA sequences of the P1 and P2 was conducted as described previously with modifications [72]. The first 25 μl PCR reaction was modified as follows: master mix containing 1.25 units of Taq DNA polymerase (iNtRON Biotechnology Inc, Gyeonggi-do, South Korea), 1х MgCl2 free PCR buffer, 2.5 mM of MgCl2, 0.2 mM of dNTP (iNtRON Biotechnology Inc, Gyeonggi-do, South Korea), 5 μl of 5 M of Betaine (Bio Basic USA Inc., New York, US) and each reaction consist of 0.5 pmol each of forward (rrs-outer-F: 5’-CTCAGAACTAACGCTGGCGGCGCG-3’), reverse primers (rrs-outer-R: 5’-GGTTCGTTACTGAGGGTTAAAACCCCC-3’) and 5 µL of the DNA from the animal urine sample. The assay was performed using the MJ Research PCT-200 Thermal Cycler (Bio-Rad, CA, USA) with the following cycling conditions: 95°C for 2 minutes; 40 cycles of 95°C for 10 seconds, 67°C for 15 seconds and 72°C for 30 seconds followed by a final extension at 72°C for 7 minutes. The second 25µl PCR reaction used 2 µL of the first PCR products in the same master mix as the first PCR, but containing 0.5 pmol each of inner forward (rrs-inner-F: 5’-CTGGCGGCGCGTCTTA-3’) and inner reverse primers (rrs-inner-R: 5’-GTTTTCACACCTGACTTACA-3’). The cycling conditions of the second PCR were modified from those of the first PCR by reducing the annealing temperature from 67°C to 55°C. The expected 547-bp amplicons were visualised using 1.5% agarose gel electrophoresis. Then, the PCR products were purified from agarose gel using GenepHlowTM Gel/PCR Kit (Geneaid, New Taipei City, Taiwan). It was conducted as described in the manufacturer’s protocol. The purified products would be sent to Bionics for the Sanger DNA Sequencing.

The chromatogram results were inspected and edited for consensus between the amplicon sequences from the reverse and forward sequencing primers using BioEdit Sequence Alignment Editor version 7.0.5.3. Maximum likelihood trees were reconstructed from the trimmed 443-nucleotide partial 16S rRNA gene alignments (from position 63 to 505 based on *L. alexanderi* GenBank accession number: NR_043047.1) on the General Time Reverse model to infer species and genetic relatedness using the Molecular Evolutionary Genetics Analysis (MEGA) software version 11 [73]. The number of base differences per sequence from averaging over all sequence pairs within each group was estimated. All sequences from this study (accession numbers: OQ446624-OQ446662)(Supplementary Table 1) and the 57 reference sequences of 40 *Leptospira* spp. acquired from GenBank data used for the analysis, (Supplementary Table 2). The initial trees for the heuristic search were obtained automatically by applying Neighbor-Join and BioNJ algorithms to a matrix of pairwise distances estimated using the Maximum Composite Likelihood (MCL) approach and selecting the topology with superior log likelihood value. A discrete Gamma distribution was used to model evolutionary rate differences among sites (5 categories, with the +G parameter set to 0.1284). The rate variation model allowed some sites to be evolutionarily invariable ([+I], 42.89% sites). The tree was drawn to scale, with branch lengths measured in the number of substitutions per site. All positions with less than 95% site coverage were eliminated, and ambiguous bases were allowed at any position (partial deletion option). The phylogenetic tree was displayed and annotated using the Interactive Tree Of Life version 6.0 [74].

### Statistical analysis

The *Leptospira* infections were estimated by stratifying each animal by sex, age group, ownership, and study sites. The *Leptospira* spp. prevalences in each exposure or condition were analysed. Odds ratios (ORs) (with the small sample adjustment) and the 95% confidence intervals (normal approximation with the small sample adjustment) were calculated to determine the strength of the associations between prevalence and exposure factors using the “epitools” package for R [75]. The analysis was done using R version 4.0.3 [76]. Map and data management were performed using Microsoft Excel.

## Results

### Asymptomatic *Leptospira* infections in dogs and cats

A total of 567 animals, comprising 303 dogs (53.4%) and 264 cats (46.6%), looked and acted healthy, were recruited into the study during the neutering service. They originated from seven provinces in three regions of Thailand: the central region (Nakhon Sawan, Nakhon Pathom, and Samut Sakhon provinces), the western region (Tak, Kanchanaburi, and Prachuap Khiri Khan provinces), and the southern region (Ranong province). The frequency of the infections in this study is shown in Fig 1. The 5/7 provincial sites show infections ranging from 1% to 16% of the recruited animals.

Of 303 dogs, 34 were infected (11.2%). The females (12.8%, 23/179) were infected more than males (8.8%, 10/114), as shown in Table 1. Meanwhile, 10.0% (1/10) of the dogs missing sex records were infected. Adults (11.9%, 32/270) were more likely to be infected than juniors (6.1%, 2/33). The infection frequency of owned dogs (10.2%, 11/108) was less than that of free-roaming dogs (11.8%, 23/195). All these differences were not statistically significant. Two of the seven study sites (Nakhon Sawan and Samut Sakhon) showed no infection. While dogs in Tak (OR = 9.125, 95% CI: 2.123-89.954, *p*-value = 0.003) and Prachuap Khiri Khan (OR = 17.290, 95% CI: 4.373-133.623, *p*-value < 0.001) study sites showed significantly higher infections than those in Nakhon Pathom site (Table 1).

**Table 1.**
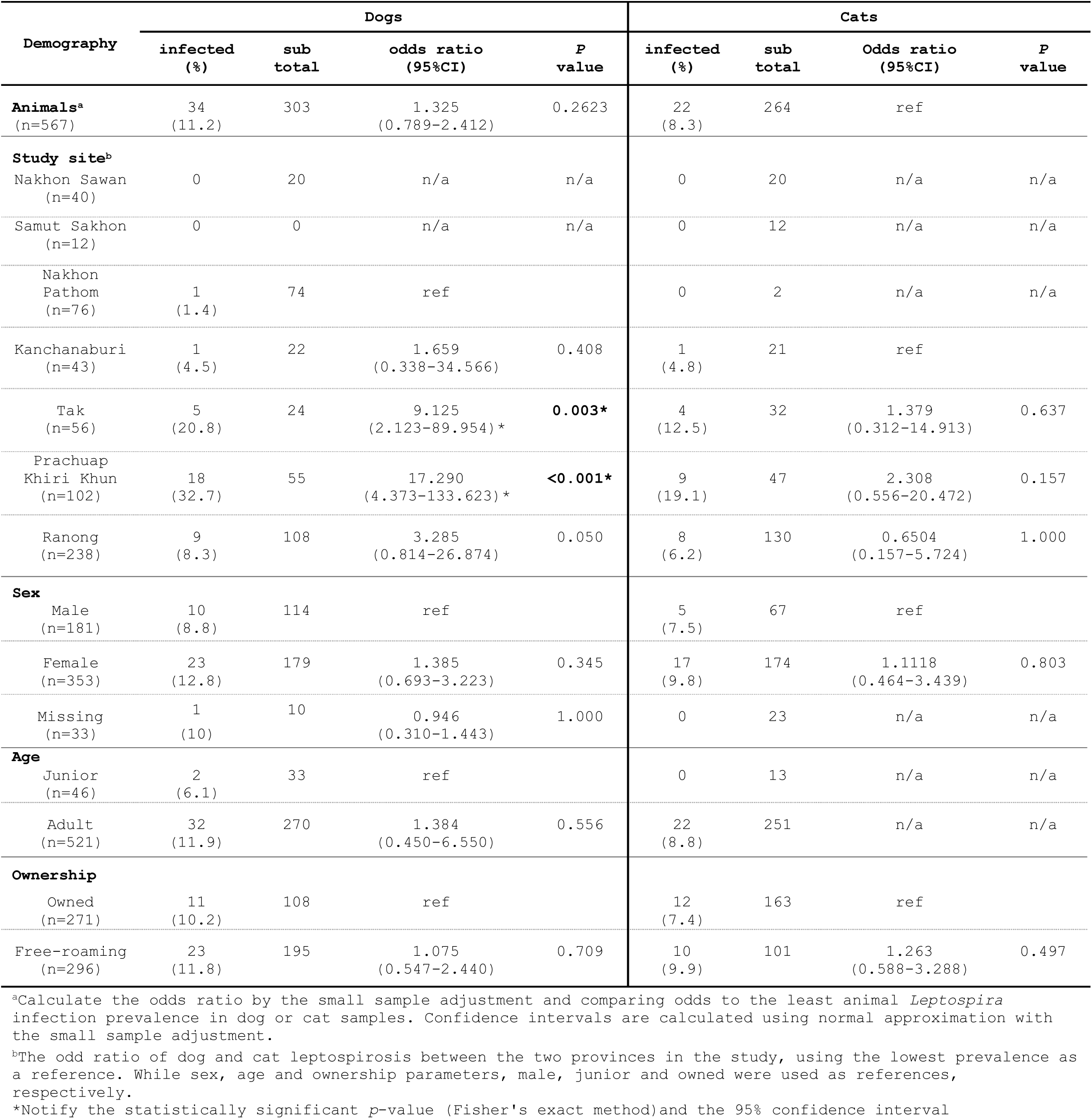
Distributions of leptospirosis in dogs and cats stratified by study sites, sex, age group and ownership.

Of the 264 cats, 22 were infected (8.3%). Like dogs, females (9.8%, 17/174) were more infected than males (7.5%, 5/67). Unlike dogs, the infections were only found in adult cats (8.8%, 22/251). The infection frequencies were observed more frequently in free-roaming cats (9.9%, 10/101) than in owned cats (7.4%, 12/163). All these differences were not statistically significant. No infections were detected in study sites located in Nakhon Sawan, Samut Sakhon, and Nakhon Pathom provinces. Among the remaining four sites, the risk of cat infections was not statistically different from that of the Kanchanaburi site (Table 1).

Human leptospirosis is a communicable disease that is part of the national surveillance program of the Department of Disease Control, Ministry of Public Health, Thailand. Suspected, probable or confirmed cases must be reported to the Department. Considering the annual incidence of human leptospirosis reported in those seven provinces (Supplementary document 1), Ranong Province showed the highest case incidence (median of 36.04 cases per 100,000 population per year) (Fig 1), followed by Tak Province (median of 1.65 cases per 100,000 population per year). The incidence in Ranong Province was about 22 times higher than in Tak Province. Compared to this study, leptospirosis in dogs and cats was more predominant in Prachuap Khiri Khan (26.5%) and Tak (16.1%) than in Ranong site studies. It is demonstrated that infections from dogs and cats can be transferred to humans. However, it was not the only source of human infections in these provinces, where other reservoirs like rodents and livestock, and environmental exposure through contaminated water or agricultural activities, also play significant roles in transmission.

### *Leptospira* Pathogen clade detected in dogs and cats

The 39/56 *Leptospira* PCR screening-positive dogs and cats had accurate partial 16S rRNA sequences (each chromatogram peak confirmed from forward and reverse sequencing primer reads). The analysis of nucleotide sequence homology and phylogenetic tree of the 443 bp of the 16S rRNA gene indicated that all were closely related to the P clade. A Maximum Likelihood phylogenetic tree was reconstructed from these 39 sequences (Supplementary Table 1) and 57 reference sequences of 40 species belonging to the P1 and P2 acquired from the GenBank database (Supplementary Table 2). Fig 2 shows that the 39 sequences from infected cats and dogs were grouped into five clusters, called after the species name of the dominating reference sequences in each cluster except the last cluster using subclade name: Interrogans (n=18, 46%), Borgpetersenii (n=5, 13%), Weilii (n=2, 5%), Yasudae (n=6, 15%), and P2 (n=8, 21%) clusters. The unknown sequences in the first three clusters were assigned species corresponding to the cluster names, and they were assigned to Group 1 of the P1 (P1-1), which are common causes of human leptospirosis worldwide. Members in these clusters were grouped with a bootstrap higher than 50% (Fig 2). The sequences of the Yasudae cluster were grouped with less than 50% bootstrap support. They were assigned to Group 2 of the P1 (P1-2), which was isolated only from environmental samples. The last cluster contained sequences from several species in the P2 (Fig 2). Within each cluster, a range of zero to eleven single-nucleotide polymorphisms (SNPs) were found (Supplementary Table 3). Of 443 bp, none of the SNPs were found within the Borgpetersenii or Weilii clusters, while six, four and eleven SNPs were found in the Interrogans, Yasudae, and P2 subclade clusters, respectively.

**Fig 2.**
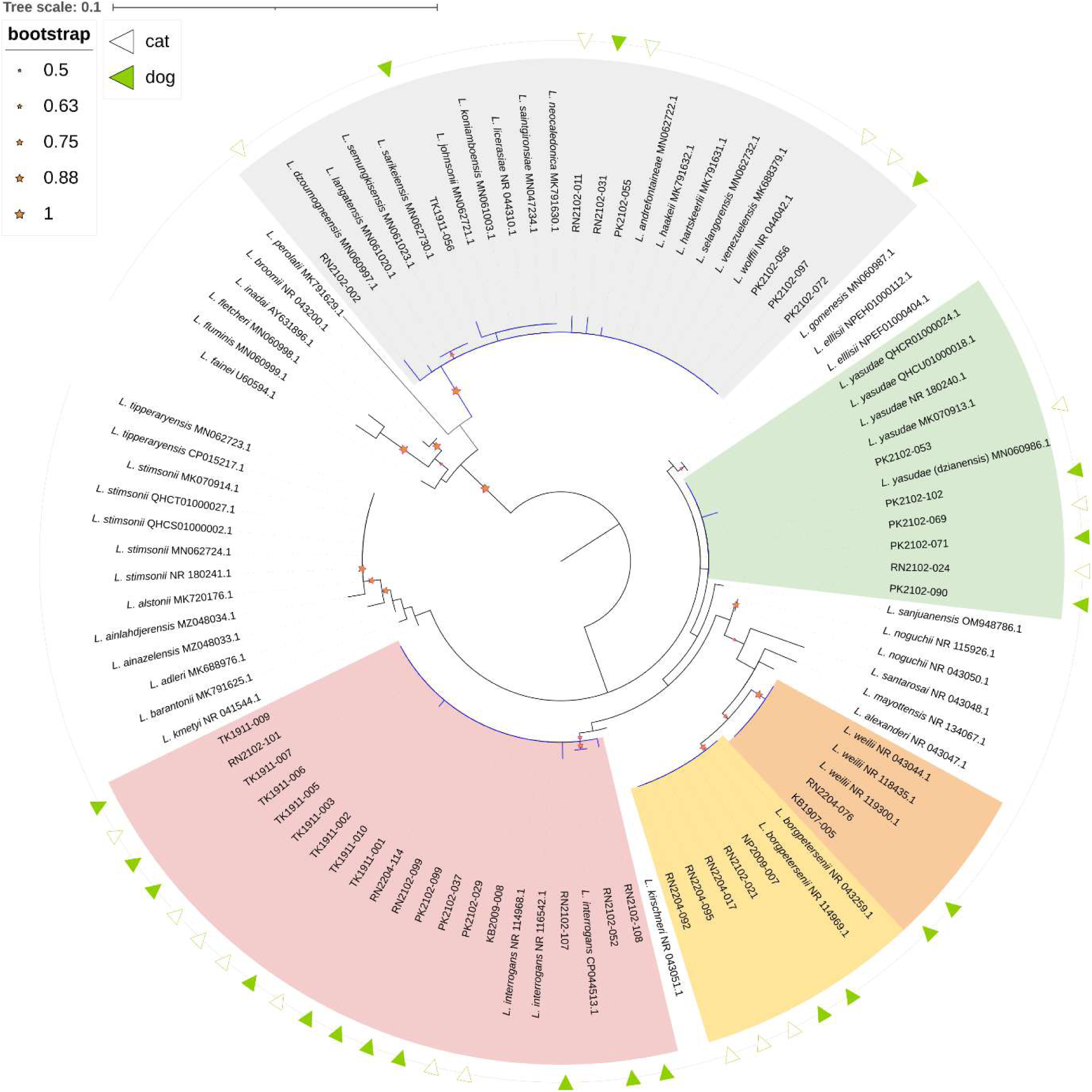
Maximum Likelihood phylogenetic tree reconstructed using the partial 16S rRNA sequences of 39 nested PCR products amplified from urine samples of dogs (filled green triangles) and cats (not filled triangles). There were 57 reference sequences of 40 *Leptospira* in the Pathogen clade, which were used to infer the species of amplicon sequences. The *Leptospira* sequences from this study were grouped into five clusters: *L. interrogans* (pink), *L. borgpetersenii* (yellow), *L. weilii* (orange), *L. yasudae* (green) and species in the Subclade 2 (grey) of the Pathogen clade. The sequences grouped in each cluster with a bootstrap value higher than 50% are indicated with a red star.

The distribution of the P clade infection in dogs and cats was summarised (Supplementary Table 4). The P1-1 infections (*L. interorgan, L. borgpetersenii* and *L. weilii*) predominated in dogs and cats originating from Tak (16.7% vs 12.5%) followed by Ranong (6.5% vs 3.1%), Kanchanaburi (4.6% vs 4.8%), Nakhon Pathom (1.4%, dogs only) and Prachuap Khiri Khun site (3.6% vs 2.1%) sites as shown in Fig 3. The *L. yasudae* (P1-2) predominated in Prachuap Khiri Khun (5.5% vs 4.3%) and Ranong (0.8%, cats only) sites, while the P2 predominated in Prachuap Khiri Khun (1.8% vs 6.4%), Ranong (0.9% vs 1.5%) and Tak sites (4.2%, dogs only).

**Fig 3.**
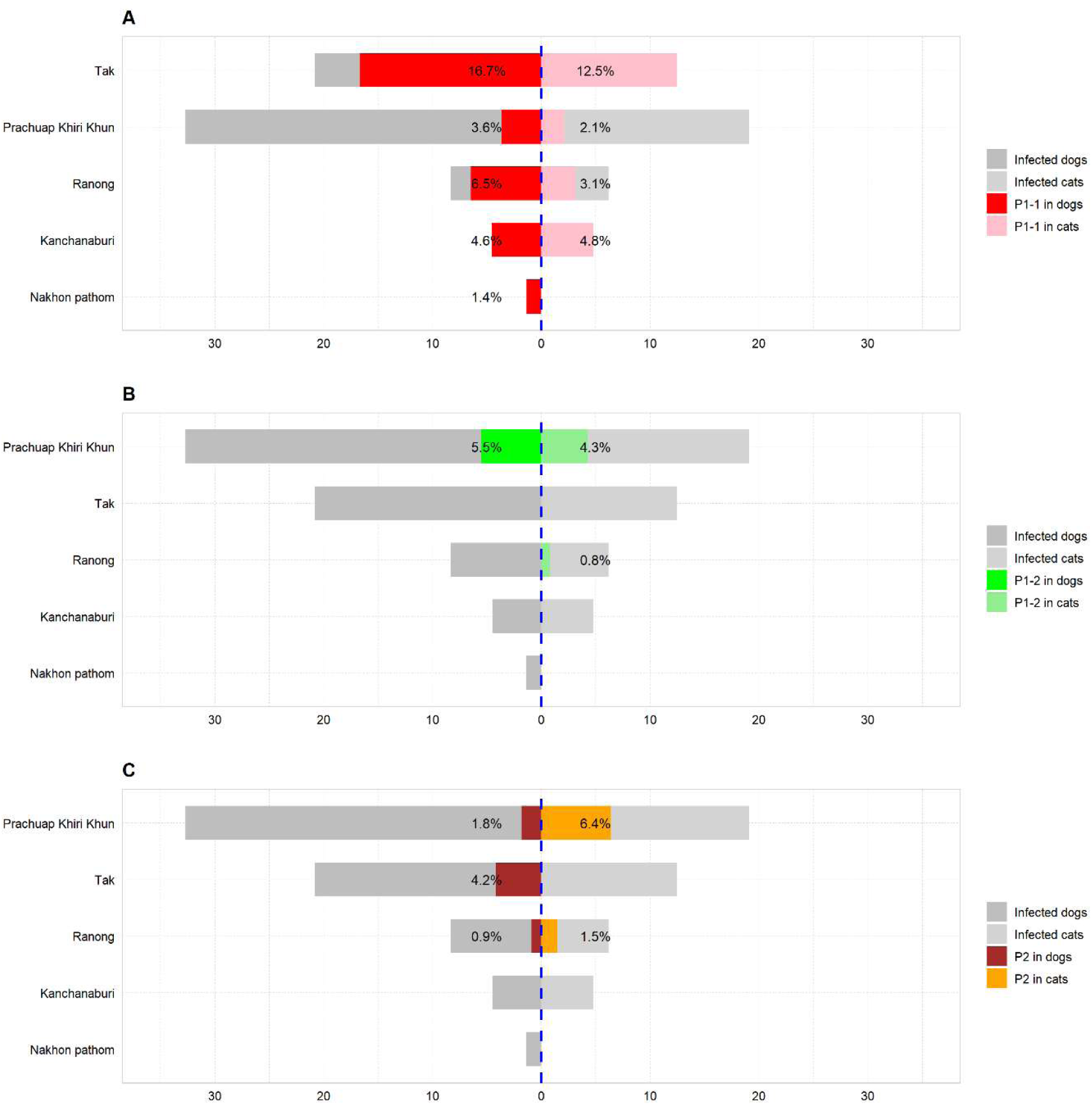
The funnel graph illustrates the distributions of Pathogen clade (P)-infected dogs (left) and cats (right) in five prevalence study sites, compared to infected animals (PCR-positive without molecular typing). *L. interrogans*, *L. borgpetersenii* and *L. weilii* (belonging to Group 1 of the Subclade 1 in the P clade as P1-1), *L. yasudae* (belonging to Group 2 of the Subclade 1 as P1-2), and species in the Subclade 2 (P2) were in Panels A, B and C, respectively.

Dogs and cats originating in Tak site were 6.122-fold more likely to be infected by the P1-1 than those in Nakhon Pathom site (95% CI: 1.498 - 51.936, *p*-value = 0.005), as shown in Table 2. Animals originating from Prachuap Khiri Khan site were 6.046-fold more likely to be infected with the P1-2 than those from Ranong site (95% CI: 1.446-55.164, *p*-value = 0.010).

**Table 2.**
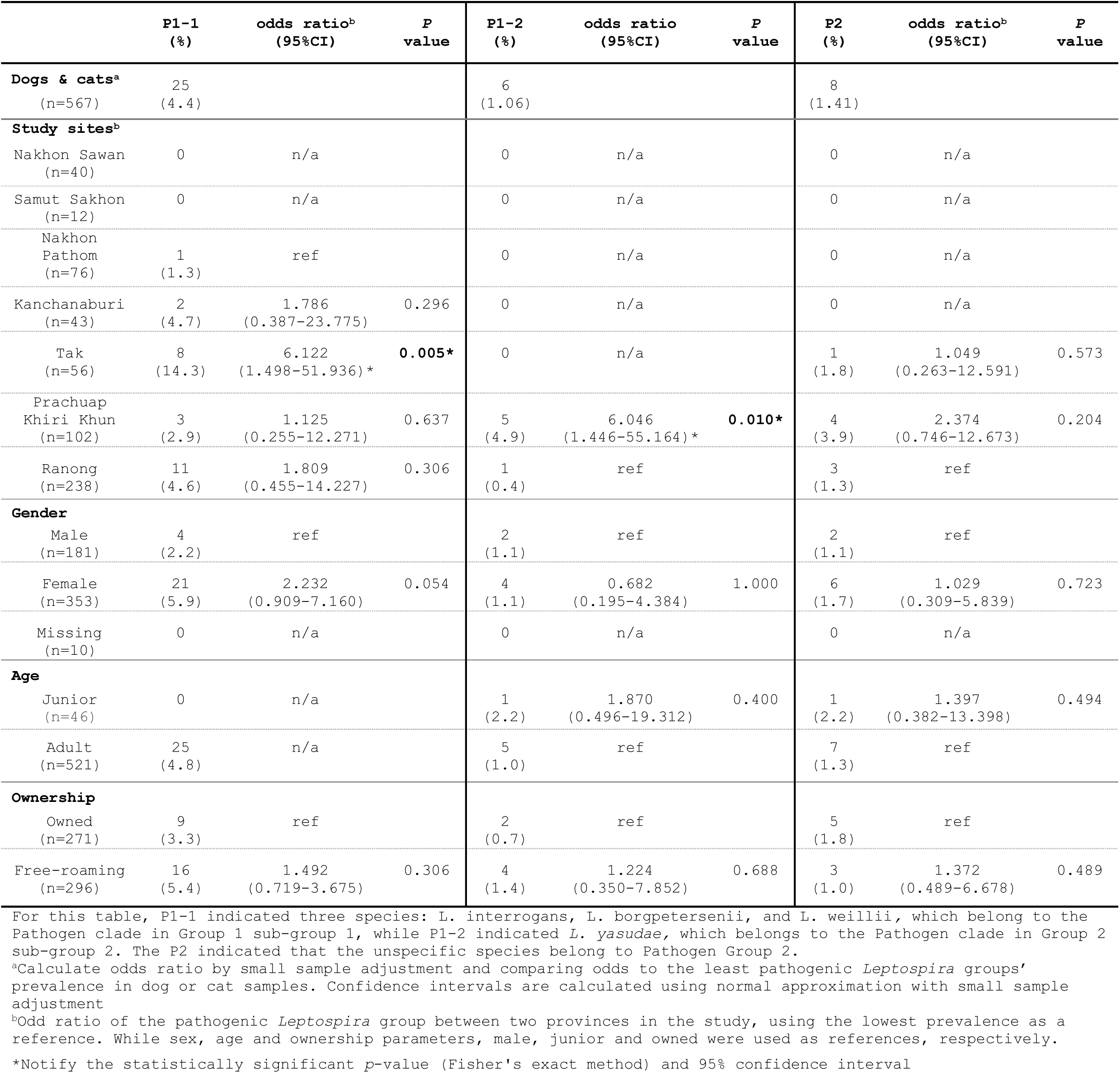
Distribution of *Leptospira* Pathogen clade detected in dogs and cats in each study site.

## Discussion

In this study, we identified urinary *Leptospira* in dogs and cats during neutering services in seven study sites in central, western and southern Thailand. *Leptospira* shedding in urine was found in 9.8% of healthy (asymptomatic infection) dogs and cats (11.2% versus 8.3% cats). *Leptospira* infections in cats and dogs were low in study sites of the central region compared to those in the western and southern regions. The prevalence of leptospirosis in dogs and cats in this study was slightly higher than the previously reported rates in Thailand, which ranged from 0.5% to 10.3% in dogs and from 0.8% to 7.8% in cats. The prevalence in dogs was comparable to those reported in Malaysia [34, 40], Germany [41], Algiers [47], Ireland [45], and the USA [38], but lower than the figures reported in Sri Lanka (15% to 15.7%) [33, 48], India (18.7%) [56], Iran (31.1%) [39], Brazil (10.6% to 19.8%) [42, 46], France (13.3%) [37], Colombia (52.9%) [55], and Ecuador (43.8% to 97.4%) [50, 53]. In cats, the prevalence fell within the previously reported range of 0.3% to 22.1%, observed in countries such as Malaysia [34, 58], Spain [65], Canada [59, 66], Germany [67], Japan [64], Chile [62], Italy [49, 61], the French Indies [63], and Brazil [69, 70]. However, strikingly higher prevalences were reported in Taiwan (67.8%) [60], Colombia (65.3%) [55], and Australia (42.4%) [68]. Variations in prevalence may be attributed to geographical location, study population, sample size, season, and screening techniques. By utilising molecular methods to specifically detect DNA of *Leptospira* species in the Pathogen Clade in this study, we demonstrate the current carrier stage among dogs and cats, which poses a potential risk of disease transmission to other mammals and humans.

Several *Leptospira* spp. within the Pathogen Clade have been reported in asymptomatic dogs and cats worldwide. These include *L. interrogans* [25, 31, 33, 34, 36, 37, 41, 42, 47–53, 55, 57, 58, 60–63, 68], *L. borgpetersenii* [33, 41, 61, 64], *L. weilii* [31, 33], *L. kirschneri* [52, 53, 60], *L. santarosai* [42, 50, 54, 57], and *L. noguchii* [25, 50]. *L. interrogans* predominates among dogs from Malaysia [34], Algiers [47], Sri Lanka [33], Brazil [25, 42, 52], New Caledonia [36], France [37], Colombia [55], Australia [68] and among cats from Malaysia [58], Taiwan [60], Chile [62], Canada [59], the French Indies [63] and Australia [68]. This study confirms that dogs and cats from central, western and southern Thailand carry these pathogenic species. We identified *L. interrogans, L. borgpetersenii*, *and L. weillii* (P1-1), which are the predominant species causing human leptospirosis in Thailand and neighbouring countries [77–84], in dogs and cats during neutering procedures in western (Tak, Kanchanaburi and Prachuap Khiri Khan), southern (Ranong) and central (Nakhon Pathom) regions of Thailand. Furthermore, this study identified unspecified species of *Leptospira* within the P2 subclade (causing mild diseases in humans [17]) and the unknown virulent *L. yasudae* (belonging to the P1-2). The P2 subclade was identified in southern (Ranong) and western (Tak and Prachuap Khiri Khan) study sites, whereas *L. yasudae* was also identified in the same regions but at the Tak site.

In this study, *L. interrogans* was identified in similar proportions in both dogs (3.6%) and cats (2.7%) from urine samples. However, a previous study reported a bias in the prevalence of *L. interrogans* infection towards cats (0.5% of dogs’ vs 7.8% of cats’ blood PCR positive) in Songkhla, southern Thailand [43]. Additionally, the prevalence rate of *L. interrogans* among dogs in this study was lower than those previously reported in Nan, northern Thailand (positive dogs’ urine at 6.9%) [31]. Nan and Songkhla were provinces, where human leptospirosis was a significant problem among the population during the wet season [21, 85]. The reason for the host bias in these study areas was unknown.

*L. borgpetersenii* is a common cause of bovine leptospirosis. The first isolates of this species were obtained from cattle at slaughter in the United States of America [86]. However, *L. borgpetersenii* infections have also been identified in dogs and cats. Notably, the number reported among cats was higher than that among dogs. For example, 7.1% of cats were infected on Okinawa Island, Japan [64], and 4.5% in southern Italy [61]. In contrast, 2.1% of dogs were infected in Sri Lanka [33] and 0.5% in Germany [41]. To our knowledge, the identification of *L. borgpetersenii* infections among asymptomatic dogs and cats has been reported for the first time in Thailand: in central (Nakhon Pathom) and southern (Ranong) regions.

*L. weilii* infections were found in several animals, including cattle, swine, canines, goat, bandicoots, and grey kangaroos [31, 87, 88]. In Thailand, canine leptospirosis was first reported in Nan, northern Thailand, with a prevalence rate of 3.5% [31]. Meanwhile, the infections in Sri Lanka demonstrated a similar prevalence of 2.1% [33]. In this study, *L. weilii* infections were found in dogs with a low prevalence of 0.7% (Kanchanaburi and Ranong) and were not identified in cats.

*L. yasudae*, firstly isolated from environmental samples, has been recently confirmed to be conspecific with *L. dzianensis* based on average nucleotide identity [7]. It was grouped into low-virulent or asymptomatic infections. There have been no reports of human or animal infection. In this study, we found that it predominantly originated in the urine samples of dogs and cats during neutering at the Prachuap Khiri Khun site, as well as in one sample from the cat at the Ranong site.

Infections of the P2 subclade were identified. However, the *rrs* sequence analysis cannot distinguish at the species level because of the limited diversity of the selected sequences among the P2 species members. Based on the single polymorphism nucleotide (SNP), three groups can be observed (Supplementary Table 3): 1) *L. andrefontaineae*, *L. haakeii*, *L. hartskeerlii*, *L. selangorensis*, *L. venezuelensis*, and *L. wolffii*; 2) *L. sarikeiensis*, *L. semungkisensis*, and *L. langatensis*; and 3) *L. neocaledonica*, *L. saintgironsiae*, *L. koniamboensis*, *L. johnsonii*, and *L. licerasiae*. Of eight unspecified species sequences of P2 subclade, six (two dogs and four cat from Ranong and Prachuap Khiri Khun sites) were closely related to the first reference group (the *L. wolffii* group) and the remaining two sequences of (one dog and one cat from Tak and Ranong sites) were closely related to the last reference group (the *L. licerasiae* group). The potential pathogenic *L. wolffii* was first identified through the isolation of a urine sample from a Thai patient with mild leptospirosis symptoms in northeastern Thailand, and it was later isolated from rodent (*Bandicota indica*) specimens in the same region [17, 89]. In Iran, the circulation of *L. wolffii* among humans and animals was reported [71, 90]. *L. wolffii* also became a major species causing human leptospirosis in north-central Bangladesh, replacing *L. interrogans* [91, 92]. These emphasise emerging *L. wolffii* infections in humans and mammals in South and Southeast Asia. While *L. licerasiae* was first discovered in rats and symptomatic human infections, which originated in or returned from South America [93, 94], more reports have been made in other regions of the world, such as clinically healthy experimentally infected Australian swine [95] and leptospirosis-vaccinated dogs in Sri Lanka [48]. The other P2 species members have been mostly isolated from the environment worldwide, including Mayotte [12], Puerto Rico [96], Japan [82, 97] and also Thailand [98]. Although we are uncertain about the true species of the P2 that infected the dogs and cats in our study, the distribution of these infections in animal reservoirs across various geographical locations should be considered in terms of their implications for animal health.

This study has limitations. Firstly, the sample sizes of dogs and cats were not equally distributed across all sites (median = 41.5, min = 12, max = 238, IQR = 56 - 89). This resulted in the underestimated infections in those low-recruitment sites. Secondly, this study was unable to present the situation of the infection throughout Thailand, as we did not investigate in northern, eastern and northeastern Thailand. Thirdly, only a proportion of positive-PCR samples were further subjected to species identification due to the limitation of amplicon sequencing, depending on the amount of *Leptospira* DNA prepared from each urine sample. Moreover, primers used in species identification may not bind efficiently to all species within the *Leptospira* pathogen clade, as we were unable to obtain high-quality forward and reverse chromatogram reads from the Sanger sequencing (which were discarded from the analysis). Due to the presence of more than one *Leptospira* infection, direct sequencing of amplified PCR products can yield ambiguous sequences or high background noise. Cloning of PCR products before DNA sequencing or next-generation deep sequencing can improve the quality of sequencing. Lastly, even though the conserved 443 bp of pathogenic *Leptospira rrs* gene sequencing analysis is a rapid method to speciate major pathogenic *Leptospira* species causing human leptospirosis (the P1 subgroup 1), it cannot accurately refer to the species within the P2 subclade, which shows a low level of genetic diversity [9]. To facilitate species identification among members of the P2 subclade, additional nucleotide sequences, adjusted to the diverse region of the P2 *rrs* gene or using other *Leptospira* diverse genes, such as *lipL32* and *secY* [96], would enable more accurate phylogenetic analysis.

In conclusion, three classical *Leptospira* species (*L. interrogans*, *L. weilii*, and *L. borgpetersenii*), which cause human leptospirosis, were predominantly carried in their urinary bladders by dogs and cats in western and southern Thailand. While other species, which are potentially causing mild leptospirosis (the *L. wolffii*-related species and the *L. licerasiae-* related species) as well as those with unknown infectivity (*L. yasudae*), were also identified. The carriage of diverse *Leptospira* species identified here emphasises the risk of leptospirosis transmission from dogs and cats to their human companions and other mammals through urination in habitats and areas where they wander. Leptospirosis vaccination, which covers a broader range of species, not just a few serovars, in animals, is considered a preventive measure against the disease and may also reduce urinary shedding of the bacteria. Our findings aim to raise public awareness of human leptospirosis transmission through contact with dogs and cats. And expand current knowledge on the epidemiology of leptospirosis and disease control in endemic areas.

## Acknowledgement

We are grateful to the staff of the One Health mobile unit, Faculty of Veterinary Science, Mahidol University, and all the volunteers from Soi Dog and the nonprofit organisations for their assistance with sample collection during the neutering process. We would also like to thank all the dog and cat owners who participated in this study.

## References

1. (WHO). WHO. Neglected tropical diseases, hidden successes, emerging opportunities Geneva: WHO Press; 2009 [11 December 2022]. Available from: https://www.who.int/publications/i/item/9789241598705.

2. Karpagam KB, Ganesh B. Leptospirosis: a neglected tropical zoonotic infection of public health importance-an updated review. Eur J Clin Microbiol Infect Dis. 2020;39(5):835–46. Epub 20200102. doi: 10.1007/s10096-019-03797-4. PubMed PMID: 31898795.

3. Adler B, de la Peña Moctezuma A. Leptospira and leptospirosis. Vet Microbiol. 2010;140(3-4):287-96. Epub 20090313. doi: 10.1016/j.vetmic.2009.03.012. PubMed PMID: 19345023.

4. Haake DA, Levett PN. Leptospirosis in humans. Curr Top Microbiol Immunol. 2015;387:65–97. doi: 10.1007/978-3-662-45059-8_5. PubMed PMID: 25388133; PubMed Central PMCID: PMCPMC4442676.

5. Evangelista KV, Coburn J. Leptospira as an emerging pathogen: a review of its biology, pathogenesis and host immune responses. Future Microbiol. 2010;5(9):1413–25. doi: 10.2217/fmb.10.102. PubMed PMID: 20860485; PubMed Central PMCID: PMCPMC3037011.

6. Casanovas-Massana A, Hamond C, Santos LA, de Oliveira D, Hacker KP, Balassiano I, et al. Leptospira yasudae sp. nov. and Leptospira stimsonii sp. nov., two new species of the pathogenic group isolated from environmental sources. Int J Syst Evol Microbiol. 2020;70(3):1450–6. doi: 10.1099/ijsem.0.003480. PubMed PMID: 31184568; PubMed Central PMCID: PMCPMC10197099.

7. Casanovas-Massana A, Vincent AT, Bourhy P, Neela VK, Veyrier FJ, Picardeau M, et al. Leptospira dzianensis and Leptospira putramalaysiae are later heterotypic synonyms of Leptospira yasudae and Leptospira stimsonii. Int J Syst Evol Microbiol. 2019;71(3). Epub 20210223. doi: 10.1099/ijsem.0.004713. PubMed PMID: 33620308; PubMed Central PMCID: PMCPMC8375420.

8. Fernandes LGV, Stone NE, Roe CC, Goris MGA, van der Linden H, Sahl JW, et al. Leptospira sanjuanensis sp. nov., a pathogenic species of the genus Leptospira isolated from soil in Puerto Rico. Int J Syst Evol Microbiol. 2022;72(10). doi: 10.1099/ijsem.0.005560. PubMed PMID: 36260655.

9. Guglielmini J, Bourhy P, Schiettekatte O, Zinini F, Brisse S, Picardeau M. Genus-wide Leptospira core genome multilocus sequence typing for strain taxonomy and global surveillance. PLoS Negl Trop Dis. 2019;13(4):e0007374. Epub 20190426. doi: 10.1371/journal.pntd.0007374. PubMed PMID: 31026256; PubMed Central PMCID: PMCPMC6513109.

10. Korba AA, Lounici H, Kainiu M, Vincent AT, Mariet JF, Veyrier FJ, et al. Leptospira ainlahdjerensis sp. nov., Leptospira ainazelensis sp. nov., Leptospira abararensis sp. nov. and Leptospira chreensis sp. nov., four new species isolated from water sources in Algeria. Int J Syst Evol Microbiol. 2021;71(12). doi: 10.1099/ijsem.0.005148. PubMed PMID: 34914572.

11. Thibeaux R, Girault D, Bierque E, Soupé-Gilbert ME, Rettinger A, Douyère A, et al. Biodiversity of Environmental Leptospira: Improving Identification and Revisiting the Diagnosis. Front Microbiol. 2018;9:816. Epub 20180501. doi: 10.3389/fmicb.2018.00816. PubMed PMID: 29765361; PubMed Central PMCID: PMCPMC5938396.

12. Vincent AT, Schiettekatte O, Goarant C, Neela VK, Bernet E, Thibeaux R, et al. Revisiting the taxonomy and evolution of pathogenicity of the genus Leptospira through the prism of genomics. PLoS Negl Trop Dis. 2019;13(5):e0007270. Epub 20190523. doi: 10.1371/journal.pntd.0007270. PubMed PMID: 31120895; PubMed Central PMCID: PMCPMC6532842.

13. Abd Rahman AN, Hasnul Hadi NH, Sun Z, Thilakavathy K, Joseph N. Regional Prevalence of Intermediate Leptospira spp. in Humans: A Meta-Analysis. Pathogens. 2021;10(8). Epub 20210727. doi: 10.3390/pathogens10080943. PubMed PMID: 34451407; PubMed Central PMCID: PMCPMC8398916.

14. Chiani Y, Jacob P, Varni V, Landolt N, Schmeling MF, Pujato N, et al. Isolation and clinical sample typing of human leptospirosis cases in Argentina. Infect Genet Evol. 2016;37:245–51. Epub 20151130. doi: 10.1016/j.meegid.2015.11.033. PubMed PMID: 26658064.

15. Chiriboga J, Barragan V, Arroyo G, Sosa A, Birdsell DN, España K, et al. High Prevalence of Intermediate Leptospira spp. DNA in Febrile Humans from Urban and Rural Ecuador. Emerg Infect Dis. 2015;21(12):2141–7. doi: 10.3201/eid2112.140659. PubMed PMID: 26583534; PubMed Central PMCID: PMCPMC4672404.

16. Perolat P, Chappel RJ, Adler B, Baranton G, Bulach DM, Billinghurst ML, et al. Leptospira fainei sp. nov., isolated from pigs in Australia. Int J Syst Bacteriol. 1998;48 Pt 3:851–8. doi: 10.1099/00207713-48-3-851. PubMed PMID: 9734039.

17. Slack AT, Kalambaheti T, Symonds ML, Dohnt MF, Galloway RL, Steigerwalt AG, et al. Leptospira wolffii sp. nov., isolated from a human with suspected leptospirosis in Thailand. Int J Syst Evol Microbiol. 2008;58(Pt 10):2305–8. doi: 10.1099/ijs.0.64947-0. PubMed PMID: 18842846.

18. Costa F, Hagan JE, Calcagno J, Kane M, Torgerson P, Martinez-Silveira MS, et al. Global Morbidity and Mortality of Leptospirosis: A Systematic Review. PLoS Negl Trop Dis. 2015;9(9):e0003898. Epub 20150917. doi: 10.1371/journal.pntd.0003898. PubMed PMID: 26379143; PubMed Central PMCID: PMCPMC4574773.

19. Victoriano AF, Smythe LD, Gloriani-Barzaga N, Cavinta LL, Kasai T, Limpakarnjanarat K, et al. Leptospirosis in the Asia Pacific region. BMC Infect Dis. 2009;9:147. Epub 20090904. doi: 10.1186/1471-2334-9-147. PubMed PMID: 19732423; PubMed Central PMCID: PMCPMC2749047.

20. Health Information System Development Office (HISO): Department of Disease Control MoPH. Thai Health Stat: Health Information System Development Office (HISO); 2020 [4 January 2024]. Available from: https://www.hiso.or.th/thaihealthstat/area/index.php?ma=1&pf=01818101&tm=2&tp=243.

21. Chadsuthi S, Chalvet-Monfray K, Geawduanglek S, Wongnak P, Cappelle J. Spatial-temporal patterns and risk factors for human leptospirosis in Thailand, 2012-2018. Sci Rep. 2022;12(1):5066. Epub 20220324. doi: 10.1038/s41598-022-09079-y. PubMed PMID: 35332199; PubMed Central PMCID: PMCPMC8948194.

22. Narkkul U, Thaipadungpanit J, Srisawat N, Rudge JW, Thongdee M, Pawarana R, et al. Human, animal, water source interactions and leptospirosis in Thailand. Sci Rep. 2021;11(1):3215. Epub 20210205. doi: 10.1038/s41598-021-82290-5. PubMed PMID: 33547388; PubMed Central PMCID: PMCPMC7864926.

23. Phosri A. Effects of rainfall on human leptospirosis in Thailand: evidence of multi-province study using distributed lag non-linear model. Stoch Environ Res Risk Assess. 2022;36(12):4119–32. Epub 20220604. doi: 10.1007/s00477-022-02250-x. PubMed PMID: 35692716; PubMed Central PMCID: PMCPMC9167037.

24. Almeida DS, Paz LN, de Oliveira DS, Silva DN, Ristow P, Hamond C, et al. Investigation of chronic infection by Leptospira spp. in asymptomatic sheep slaughtered in slaughterhouse. PLoS One. 2019;14(5):e0217391. Epub 20190523. doi: 10.1371/journal.pone.0217391. PubMed PMID: 31120961; PubMed Central PMCID: PMCPMC6532964.

25. Sant’Anna da Costa R,> MI NDA, Dos Santos Baptista Borges AL, Carvalho-Costa FA, Martins G, Lilenbaum W. Persistent High Leptospiral Shedding by Asymptomatic Dogs in Endemic Areas Triggers a Serious Public Health Concern. Animals (Basel). 2021;11(4). Epub 20210326. doi: 10.3390/ani11040937. PubMed PMID: 33810226; PubMed Central PMCID: PMCPMC8065945.

26. T. V. Pet parenting on the rise Bangkok: Thai Public Broadcasting Service; 2021 [cited 2024 4 January]. Available from: https://www.thaipbsworld.com/pet-parenting-on-the-rise/.

27. Puranabhandu O ME. Thailand’s Pet Food Market Bangkok: The United States Department of Agriculture, Foreign Agricultural Service; 2021 [4 January 2024]. Available from: https://fas.usda.gov/data/thailand-thailands-pet-food-market.

28. Srimoragot P SP, Suwannaphirom P, Ruchisereekul K. Surveillance of knowledge, and opinion among pet’s owner on tobacco effect on pets’ health. J Mahanakorn Vet Med. 2021;16(1):63-75.

29. Thanapongtharm W, Kasemsuwan S, Wongphruksasoong V, Boonyo K, Pinyopummintr T, Wiratsudakul A, et al. Spatial Distribution and Population Estimation of Dogs in Thailand: Implications for Rabies Prevention and Control. Front Vet Sci. 2021;8:790701. Epub 20211221. doi: 10.3389/fvets.2021.790701. PubMed PMID: 34993247; PubMed Central PMCID: PMCPMC8724437.

30. Altheimer K, Jongwattanapisan P, Luengyosluechakul S, Pusoonthornthum R, Prapasarakul N, Kurilung A, et al. Leptospira infection and shedding in dogs in Thailand. BMC Vet Res. 2020;16(1):89. Epub 20200317. doi: 10.1186/s12917-020-2230-0. PubMed PMID: 32178664; PubMed Central PMCID: PMCPMC7077098.

31. Kurilung A, Chanchaithong P, Lugsomya K, Niyomtham W, Wuthiekanun V, Prapasarakul N. Molecular detection and isolation of pathogenic Leptospira from asymptomatic humans, domestic animals and water sources in Nan province, a rural area of Thailand. Res Vet Sci. 2017;115:146–54. Epub 20170329. doi: 10.1016/j.rvsc.2017.03.017. PubMed PMID: 28384550.

32. Sprißler F, Jongwattanapisan P, Luengyosluechakul S, Pusoonthornthum R, Prapasarakul N, Kurilung A, et al. Leptospira infection and shedding in cats in Thailand. Transbound Emerg Dis. 2019;66(2):948–56. Epub 20190107. doi: 10.1111/tbed.13110. PubMed PMID: 30580489.

33. Athapattu T, Fernando R, Abayawansha R, Fernando P, Fuward M, Samarakoon N, et al. Carrier Status of Leptospira spp. in Healthy Companion Dogs in Sri Lanka. Vector Borne Zoonotic Dis. 2022;22(2):93–100. Epub 20220128. doi: 10.1089/vbz.2021.0065. PubMed PMID: 35099292.

34. Benacer D, Thong KL, Ooi PT, Souris M, Lewis JW, Ahmed AA, et al. Serological and molecular identification of Leptospira spp. in swine and stray dogs from Malaysia. Trop Biomed. 2017;34(1):89–97. PubMed PMID: 33592986.

35. Delaude A, Rodriguez-Campos S, Dreyfus A, Counotte MJ, Francey T, Schweighauser A, et al. Canine leptospirosis in Switzerland-A prospective cross-sectional study examining seroprevalence, risk factors and urinary shedding of pathogenic leptospires. Prev Vet Med. 2017;141:48–60. Epub 20170426. doi: 10.1016/j.prevetmed.2017.04.008. PubMed PMID: 28532993.

36. Gay N, Soupé-Gilbert ME, Goarant C. Though not reservoirs, dogs might transmit Leptospira in New Caledonia. Int J Environ Res Public Health. 2014;11(4):4316–25. Epub 20140417. doi: 10.3390/ijerph110404316. PubMed PMID: 24747539; PubMed Central PMCID: PMCPMC4025015.

37. Goy-Thollot I, Djelouadji Z, Nennig M, Hazart G, Hugonnard M. Screening for Leptospira DNA in blood and urine from 30 apparently healthy dogs. Revue Vétérinaire Clinique. 2018;53(3):79–86. doi: 10.1016/j.anicom.2018.06.003.

38. Harkin KR, Roshto YM, Sullivan JT, Purvis TJ, Chengappa MM. Comparison of polymerase chain reaction assay, bacteriologic culture, and serologic testing in assessment of prevalence of urinary shedding of leptospires in dogs. J Am Vet Med Assoc. 2003;222(9):1230–3. doi: 10.2460/javma.2003.222.1230. PubMed PMID: 12725310.

39. Khorami N, Malmasi A, Zakeri S, Zahraei Salehi T, Abdollahpour G, Nassiri SM, et al. Screening urinalysis in dogs with urinary shedding of leptospires. Comparative Clinical Pathology. 2010;19(3):271–4. doi: 10.1007/s00580-009-0856-1. PubMed PMID: 1112235319.

40. Lau SF, Low KN, Khor KH, Roslan MA, Bejo SK, Radzi R, et al. Prevalence of leptospirosis in healthy dogs and dogs with kidney disease in Klang Valley, Malaysia. Trop Biomed. 2016;33(3):469–75. PubMed PMID: 33579118.

41. Llewellyn JR, Krupka-Dyachenko I, Rettinger AL, Dyachenko V, Stamm I, Kopp PA, et al. Urinary shedding of leptospires and presence of Leptospira antibodies in healthy dogs from Upper Bavaria. Berl Munch Tierarztl Wochenschr. 2016;129(5-6):251–7. PubMed PMID: 27344919.

42. Miotto BA, Guilloux AGA, Tozzi BF, Moreno LZ, da Hora AS, Dias RA, et al. Prospective study of canine leptospirosis in shelter and stray dog populations: Identification of chronic carriers and different Leptospira species infecting dogs. PLoS One. 2018;13(7):e0200384. Epub 20180711. doi: 10.1371/journal.pone.0200384. PubMed PMID: 29995963; PubMed Central PMCID: PMCPMC6040711.

43. Ngasaman R, Saechan V, Prachantasena S, Yingkajorn M, Sretrirutchai S. Investigation of Leptospira Infection in Stray Animals in Songkhla, Thailand: Leptospirosis Risk Reduction in Human. Vector Borne Zoonotic Dis. 2020;20(6):432–5. Epub 20200106. doi: 10.1089/vbz.2019.2549. PubMed PMID: 31905047.

44. Rohilla P, Khurana R, Kumar A, Batra K, Gupta R. Detection of Leptospira in urine of apparently healthy dogs by quantitative polymerase chain reaction in Haryana, India. Vet World. 2020;13(11):2411–5. Epub 20201112. doi: 10.14202/vetworld.2020.2411-2415. PubMed PMID: 33363334; PubMed Central PMCID: PMCPMC7750226.

45. Rojas P, Monahan AM, Schuller S, Miller IS, Markey BK, Nally JE. Detection and quantification of leptospires in urine of dogs: a maintenance host for the zoonotic disease leptospirosis. Eur J Clin Microbiol Infect Dis. 2010;29(10):1305–9. Epub 20100618. doi: 10.1007/s10096-010-0991-2. PubMed PMID: 20559675.

46. Sant’anna R, Vieira AS, Grapiglia J, Lilenbaum W. High number of asymptomatic dogs as leptospiral carriers in an endemic area indicates a serious public health concern. Epidemiol Infect. 2017;145(9):1852–4. Epub 20170403. doi: 10.1017/s0950268817000632. PubMed PMID: 28367783; PubMed Central PMCID: PMCPMC9203335.

47. Zaidi S, Bouam A, Bessas A, Hezil D, Ghaoui H, Ait-Oudhia K, et al. Urinary shedding of pathogenic Leptospira in stray dogs and cats, Algiers: A prospective study. PLoS One. 2018;13(5):e0197068. Epub 20180516. doi: 10.1371/journal.pone.0197068. PubMed PMID: 29768448; PubMed Central PMCID: PMCPMC5955589.

48. Gamage CD, Sykes JE, Athapattu TPJ, Senerathne P, Karunadasa U, Fuward M, et al. Isolation of Leptospira licerasiae, Leptospira interrogans and Leptospira kmetyi From Apparently Healthy Companion Dogs Vaccinated for Leptospirosis. Vet Med Sci. 2025;11(3):e70375. doi: 10.1002/vms3.70375. PubMed PMID: 40309759; PubMed Central PMCID: PMCPMC12044410.

49. Grippi F, Cannella V, Macaluso G, Blanda V, Emmolo G, Santangelo F, et al. Serological and Molecular Evidence of Pathogenic Leptospira spp. in Stray Dogs and Cats of Sicily (South Italy), 2017-2021. Microorganisms. 2023;11(2). Epub 20230202. doi: 10.3390/microorganisms11020385. PubMed PMID: 36838350; PubMed Central PMCID: PMCPMC9963455.

50. Guzmán DA, Diaz E, Sáenz C, Álvarez H, Cueva R, Zapata-Ríos G, et al. Domestic dogs in indigenous Amazonian communities: Key players in Leptospira cycling and transmission? PLoS Negl Trop Dis. 2024;18(4):e0011671. Epub 20240403. doi: 10.1371/journal.pntd.0011671. PubMed PMID: 38568912; PubMed Central PMCID: PMCPMC10990217.

51. Mazzotta E, Lucchese L, Corrò M, Ceglie L, Danesi P, Capello K, et al. Zoonoses in dog and cat shelters in North-East Italy: update on emerging, neglected and known zoonotic agents. Front Vet Sci. 2024;11:1490649. Epub 20241127. doi: 10.3389/fvets.2024.1490649. PubMed PMID: 39664895; PubMed Central PMCID: PMCPMC11631924.

52. Merker Breyer G, Noronha Arechavaleta N, Corrêa da Silva B, Rocha Jacques da Silva ME, Costa Torres M, Cadó Nemitz L, et al. Canine Leptospirosis in Flood-Affected Areas of Southern Brazil: Molecular Assessment and Public Health Implications. Infect Dis Rep. 2025;17(3). Epub 20250603. doi: 10.3390/idr17030063. PubMed PMID: 40559194; PubMed Central PMCID: PMCPMC12193477.

53. Orlando SA, Mora-Jaramillo N, Paredes-Núñez D, Rodriguez-Pazmiño AS, Carvajal E, León Sosa A, et al. Leptospirosis outbreak in Ecuador in 2023: A pilot study for surveillance from a One Health perspective. One Health. 2024;19:100948. Epub 20241202. doi: 10.1016/j.onehlt.2024.100948. PubMed PMID: 39717537; PubMed Central PMCID: PMCPMC11664413.

54. Perez-Garcia J, Monroy FP, Agudelo-Florez P. Canine Leptospirosis in a Northwestern Region of Colombia: Serological, Molecular and Epidemiological Factors. Pathogens. 2022;11(9). Epub 20220913. doi: 10.3390/pathogens11091040. PubMed PMID: 36145472; PubMed Central PMCID: PMCPMC9506147.

55. Puentes MMM, Camargo KDJ, Roberto YAM, Guzman-Barragan BL, Tafur-Gomez GA, Clavijo NFS. Infection and re-infection of Leptospira spp. in stray dogs and cats from Bogota, Colombia. Vet World. 2024;17(5):973–80. Epub 20240504. doi: 10.14202/vetworld.2024.973-980. PubMed PMID: 38911095; PubMed Central PMCID: PMCPMC11188880.

56. Sarangi S, Vijaya Bharathi M, Madhanmohan M, Meenambigai TV, Soundararajan C, Manimaran K, et al. Molecular and serological detection of acute canine leptospirosis and associated predictive risk factors in and around Chennai, India. Microb Pathog. 2025;198:107120. Epub 20241115. doi: 10.1016/j.micpath.2024.107120. PubMed PMID: 39549929.

57. Silva-Ramos CR, Lemaitre GP, Mejorano-Fonseca JA, Matiz-González JM, Aricapa-Giraldo HJ, Agudelo JC, et al. Molecular Evidence of Leptospira spp. Infection Among Household Dogs From 15 Municipalities of the Department of Caldas, Colombia. Zoonoses Public Health. 2025;72(2):215–22. Epub 20241210. doi: 10.1111/zph.13204. PubMed PMID: 39658809.

58. Alashraf AR, Lau SF, Khairani-Bejo S, Khor KH, Ajat M, Radzi R, et al. First report of pathogenic Leptospira spp. isolated from urine and kidneys of naturally infected cats. PLoS One. 2020;15(3):e0230048. Epub 20200310. doi: 10.1371/journal.pone.0230048. PubMed PMID: 32155209; PubMed Central PMCID: PMCPMC7064249.

59. Bourassi E, Savidge C, Foley P, Hartwig S. Serologic and urinary survey of exposure to Leptospira species in a feral cat population of Prince Edward Island, Canada. J Feline Med Surg. 2021;23(12):1155–61. Epub 20210315. doi: 10.1177/1098612x211001042. PubMed PMID: 33719673; PubMed Central PMCID: PMCPMC8637349.

60. Chan KW, Hsu YH, Hu WL, Pan MJ, Lai JM, Huang KC, et al. Serological and PCR detection of feline leptospira in southern Taiwan. Vector Borne Zoonotic Dis. 2014;14(2):118–23. Epub 20131220. doi: 10.1089/vbz.2013.1324. PubMed PMID: 24359421.

61. Donato G, Masucci M, Hartmann K, Goris MGA, Ahmed AA, Archer J, et al. Leptospira spp. Prevalence in Cats from Southern Italy with Evaluation of Risk Factors for Exposure and Clinical Findings in Infected Cats. Pathogens. 2022;11(10). Epub 20220930. doi: 10.3390/pathogens11101129. PubMed PMID: 36297186; PubMed Central PMCID: PMCPMC9609655.

62. Dorsch R, Ojeda J, Salgado M, Monti G, Collado B, Tomckowiack C, et al. Cats shedding pathogenic Leptospira spp.-An underestimated zoonotic risk? PLoS One. 2020;15(10):e0239991. Epub 20201022. doi: 10.1371/journal.pone.0239991. PubMed PMID: 33091006; PubMed Central PMCID: PMCPMC7580889.

63. Holzapfel M, Taraveau F, Djelouadji Z. Serological and molecular detection of pathogenic Leptospira in domestic and stray cats on Reunion Island, French Indies. Epidemiol Infect. 2021;149:e229. Epub 20210810. doi: 10.1017/s095026882100176x. PubMed PMID: 34372952; PubMed Central PMCID: PMCPMC8569831.

64. Kakita T, Kuba Y, Kyan H, Okano S, Morita M, Koizumi N. Molecular and serological epidemiology of Leptospira infection in cats in Okinawa Island, Japan. Sci Rep. 2021;11(1):10365. Epub 20210514. doi: 10.1038/s41598-021-89872-3. PubMed PMID: 33990653; PubMed Central PMCID: PMCPMC8121857.

65. Murillo A, Cuenca R, Serrano E, Marga G, Ahmed A, Cervantes S, et al. Leptospira Detection in Cats in Spain by Serology and Molecular Techniques. Int J Environ Res Public Health. 2020;17(5). Epub 20200302. doi: 10.3390/ijerph17051600. PubMed PMID: 32121670; PubMed Central PMCID: PMCPMC7084519.

66. Rodriguez J, Blais MC, Lapointe C, Arsenault J, Carioto L, Harel J. Serologic and urinary PCR survey of leptospirosis in healthy cats and in cats with kidney disease. J Vet Intern Med. 2014;28(2):284–93. Epub 20140113. doi: 10.1111/jvim.12287. PubMed PMID: 24417764; PubMed Central PMCID: PMCPMC4858000.

67. Weis S, Rettinger A, Bergmann M, Llewellyn JR, Pantchev N, Straubinger RK, et al. Detection of Leptospira DNA in urine and presence of specific antibodies in outdoor cats in Germany. J Feline Med Surg. 2017;19(4):470–6. Epub 20160709. doi: 10.1177/1098612x16634389. PubMed PMID: 26927819.

68. Dybing NA, Jacobson C, Irwin P, Algar D, Adams PJ. Leptospira Species in Feral Cats and Black Rats from Western Australia and Christmas Island. Vector Borne Zoonotic Dis. 2017;17(5):319–24. Epub 20170306. doi: 10.1089/vbz.2016.1992. PubMed PMID: 28437186.

69. Paim MG, Rivas BB, Sebastião GA, Kaefer K, Rodrigues RO, Mayer FQ, et al. Investigation of anti-Leptospira spp. antibodies and leptospiruria in cats attended to a veterinary teaching hospital in southern Brazil. Comp Immunol Microbiol Infect Dis. 2024;107:102138. Epub 20240203. doi: 10.1016/j.cimid.2024.102138. PubMed PMID: 38367297.

70. Ulsenheimer BC, Tonin AA, von Laer AE, Dos Santos HF, Sangioni LA, Fighera R, et al. Molecular detection and phylogenetic analysis of Leptospira interrogans and Leptospira borgpetersenii in cats from Central region of Rio Grande do Sul state, Brazil. Comp Immunol Microbiol Infect Dis. 2025;116:102286. Epub 20241128. doi: 10.1016/j.cimid.2024.102286. PubMed PMID: 39644868.

71. Zakeri S, Khorami N, Ganji ZF, Sepahian N, Malmasi AA, Gouya MM, et al. Leptospira wolffii, a potential new pathogenic Leptospira species detected in human, sheep and dog. Infect Genet Evol. 2010;10(2):273–7. Epub 20100114. doi: 10.1016/j.meegid.2010.01.001. PubMed PMID: 20074666.

72. Boonsilp S, Thaipadungpanit J, Amornchai P, Wuthiekanun V, Chierakul W, Limmathurotsakul D, et al. Molecular detection and speciation of pathogenic Leptospira spp. in blood from patients with culture-negative leptospirosis. BMC Infect Dis. 2011;11:338. Epub 20111213. doi: 10.1186/1471-2334-11-338. PubMed PMID: 22151687; PubMed Central PMCID: PMCPMC3297668.

73. Tamura K SG, Kumar S. MEGA 11: Molecular Evolutionary Genetics Analysis. Version 11 [software]. Molecular Biology and Evolution; 2021.

74. Letunic I, Bork P. Interactive Tree of Life (iTOL) v6: recent updates to the phylogenetic tree display and annotation tool. Nucleic Acids Res. 2024;52(W1):W78–w82. doi: 10.1093/nar/gkae268. PubMed PMID: 38613393; PubMed Central PMCID: PMCPMC11223838.

75. Tomas J. Aragon MPF DW, Adam Omidpanah. Tools for training and practicing epidemiologists including methods for two-way and multi-way contingency tables. 2020.

76. Team RC. R: A language and environment for statistical computing. 4.0.3 ed. Vienna, Austria.: R Foundation for Statistical Computing; 2020.

77. Chutinantakul A CP, Buathong R. Outbreaks of leptospirosis after a flood in Thung Song District, Nakhon Si Thammarat, January 2017. Dis Control J. 2019;45:317–29. doi: 10.14456/dcj.2019.30.

78. Ehelepola NDB, Ariyaratne K, Dissanayake DS. The interrelationship between meteorological parameters and leptospirosis incidence in Hambantota district, Sri Lanka 2008-2017 and practical implications. PLoS One. 2021;16(1):e0245366. Epub 20210122. doi: 10.1371/journal.pone.0245366. PubMed PMID: 33481868; PubMed Central PMCID: PMCPMC7822256.

79. Hinjoy S, Kongyu S, Doung-Ngern P, Doungchawee G, Colombe SD, Tsukayama R, et al. Environmental and Behavioral Risk Factors for Severe Leptospirosis in Thailand. Trop Med Infect Dis. 2019;4(2). Epub 20190516. doi: 10.3390/tropicalmed4020079. PubMed PMID: 31100812; PubMed Central PMCID: PMCPMC6631942.

80. Kakita T, Okano S, Kyan H, Miyahira M, Taira K, Kitashoji E, et al. Laboratory diagnostic, epidemiological, and clinical characteristics of human leptospirosis in Okinawa Prefecture, Japan, 2003-2020. PLoS Negl Trop Dis. 2021;15(12):e0009993. Epub 20211214. doi: 10.1371/journal.pntd.0009993. PubMed PMID: 34905535; PubMed Central PMCID: PMCPMC8670671.

81. Philip N, Ahmed K. Leptospirosis in Malaysia: current status, insights, and future prospects. J Physiol Anthropol. 2023;42(1):30. Epub 20231212. doi: 10.1186/s40101-023-00347-y. PubMed PMID: 38087323; PubMed Central PMCID: PMCPMC10714552.

82. Sato Y, Hermawan I, Kakita T, Okano S, Imai H, Nagai H, et al. Analysis of human clinical and environmental Leptospira to elucidate the eco-epidemiology of leptospirosis in Yaeyama, subtropical Japan. PLoS Negl Trop Dis. 2022;16(3):e0010234. Epub 20220331. doi: 10.1371/journal.pntd.0010234. PubMed PMID: 35358181; PubMed Central PMCID: PMCPMC8970387.

83. Thaipadungpanit J, Wuthiekanun V, Chierakul W, Smythe LD, Petkanchanapong W, Limpaiboon R, et al. A dominant clone of Leptospira interrogans associated with an outbreak of human leptospirosis in Thailand. PLoS Negl Trop Dis. 2007;1(1):e56. Epub 20071031. doi: 10.1371/journal.pntd.0000056. PubMed PMID: 17989782; PubMed Central PMCID: PMCPMC2041815.

84. Zhang R, Zhou W, Ye Q, Song S, Wang Y, Xu Y, et al. Comparative genomic analysis of Chinese human leptospirosis vaccine strain and circulating isolate. Hum Vaccin Immunother. 2020;16(6):1345–53. Epub 20200211. doi: 10.1080/21645515.2020.1720439. PubMed PMID: 32045318; PubMed Central PMCID: PMCPMC7538020.

85. Della Rossa P, Tantrakarnapa K, Sutdan D, Kasetsinsombat K, Cosson JF, Supputamongkol Y, et al. Environmental factors and public health policy associated with human and rodent infection by leptospirosis: a land cover-based study in Nan province, Thailand. Epidemiol Infect. 2016;144(7):1550–62. Epub 20151126. doi: 10.1017/s0950268815002903. PubMed PMID: 26607833; PubMed Central PMCID: PMCPMC9150581.

86. Putz EJ, Sivasankaran SK, Fernandes LGV, Brunelle B, Lippolis JD, Alt DP, et al. Distinct transcriptional profiles of Leptospira borgpetersenii serovar Hardjo strains JB197 and HB203 cultured at different temperatures. PLoS Negl Trop Dis. 2021;15(4):e0009320. Epub 20210407. doi: 10.1371/journal.pntd.0009320. PubMed PMID: 33826628; PubMed Central PMCID: PMCPMC8055020.

87. Roberts MW, Smythe L, Dohnt M, Symonds M, Slack A. Serologic-based investigation of leptospirosis in a population of free-ranging eastern grey kangaroos (Macropus giganteus) indicating the presence of Leptospira weilii serovar Topaz. J Wildl Dis. 2010;46(2):564–9. doi: 10.7589/0090-3558-46.2.564. PubMed PMID: 20688651.

88. Slack AT, Symonds ML, Dohnt MF, Corney BG, Smythe LD. Epidemiology of Leptospira weilii serovar Topaz infections in Australia. Commun Dis Intell Q Rep. 2007;31(2):216–22. doi: 10.33321/cdi.2007.31.19. PubMed PMID: 17724998.

89. Krairojananan P, Thaipadungpanit J, Leepitakrat S, Monkanna T, Wanja EW, Schuster AL, et al. Low Prevalence of Leptospira Carriage in Rodents in Leptospirosis-Endemic Northeastern Thailand. Trop Med Infect Dis. 2020;5(4). Epub 20200930. doi: 10.3390/tropicalmed5040154. PubMed PMID: 33008058; PubMed Central PMCID: PMCPMC7720114.

90. Zakeri S, Sepahian N, Afsharpad M, Esfandiari B, Ziapour P, Djadid ND. Molecular epidemiology of leptospirosis in northern Iran by nested polymerase chain reaction/restriction fragment length polymorphism and sequencing methods. Am J Trop Med Hyg. 2010;82(5):899–903. doi: 10.4269/ajtmh.2010.09-0721. PubMed PMID: 20439973; PubMed Central PMCID: PMCPMC2861398.

91. Djadid ND, Ganji ZF, Gouya MM, Rezvani M, Zakeri S. A simple and rapid nested polymerase chain reaction-restriction fragment length polymorphism technique for differentiation of pathogenic and nonpathogenic Leptospira spp. Diagn Microbiol Infect Dis. 2009;63(3):251–6. Epub 20081219. doi: 10.1016/j.diagmicrobio.2008.10.017. PubMed PMID: 19097839.

92. Rahman S, Paul SK, Aung MS, Ahmed S, Haque N, Raisul MNI, et al. Predominance of Leptospira wolffii in north-central Bangladesh, 2019. New Microbes New Infect. 2020;38:100765. Epub 20200919. doi: 10.1016/j.nmni.2020.100765. PubMed PMID: 33133612; PubMed Central PMCID: PMCPMC7588863.

93. Matthias MA, Ricaldi JN, Cespedes M, Diaz MM, Galloway RL, Saito M, et al. Human leptospirosis caused by a new, antigenically unique Leptospira associated with a Rattus species reservoir in the Peruvian Amazon. PLoS Negl Trop Dis. 2008;2(4):e213. Epub 20080402. doi: 10.1371/journal.pntd.0000213. PubMed PMID: 18382606; PubMed Central PMCID: PMCPMC2271056.

94. Tsuboi M, Koizumi N, Hayakawa K, Kanagawa S, Ohmagari N, Kato Y. Imported Leptospira licerasiae Infection in Traveler Returning to Japan from Brazil. Emerg Infect Dis. 2017;23(3):548–9. doi: 10.3201/eid2303.161262. PubMed PMID: 28221126; PubMed Central PMCID: PMCPMC5382744.

95. Steinrigl A, Willixhofer D, Schindler M, Richter S, Unterweger C, Ahmed AA, et al. Isolation and characterization of Leptospira licerasiae in Austrian swine - a first-time case report in Europe. BMC Vet Res. 2024;20(1):348. Epub 20240807. doi: 10.1186/s12917-024-04213-6. PubMed PMID: 39113014; PubMed Central PMCID: PMCPMC11304667.

96. Stone NE, Hall CM, Ortiz M, Hutton SM, Santana-Propper E, Celona KR, et al. Diverse lineages of pathogenic Leptospira species are widespread in the environment in Puerto Rico, USA. PLoS Negl Trop Dis. 2022;16(5):e0009959. Epub 20220518. doi: 10.1371/journal.pntd.0009959. PubMed PMID: 35584143; PubMed Central PMCID: PMCPMC9154103.

97. Masuzawa T, Sakakibara K, Saito M, Hidaka Y, Villanueva S, Yanagihara Y, et al. Characterization of Leptospira species isolated from soil collected in Japan. Microbiol Immunol. 2018;62(1):55–9. Epub 20171212. doi: 10.1111/1348-0421.12551. PubMed PMID: 29105847.

98. Chaiwattanarungruengpaisan S, Thepapichaikul W, Paungpin W, Ketchim K, Suwanpakdee S, Thongdee M. Potentially Pathogenic Leptospira in the Environment of an Elephant Camp in Thailand. Trop Med Infect Dis. 2020;5(4). Epub 20201206. doi: 10.3390/tropicalmed5040183. PubMed PMID: 33291225; PubMed Central PMCID: PMCPMC7768412.

